# Assessing Attention Orienting in Mice: A Novel Touchscreen Adaptation of the Posner-Style Cueing Task

**DOI:** 10.1101/2020.06.05.136689

**Authors:** S. Li, C. May, AJ. Hannan, KA. Johnson, EL. Burrows

## Abstract

Atypical attention orienting has been found to be impaired in many neuropsychological disorders, but the underlying neural mechanism remains unclear. Attention can be oriented exogenously (i.e., driven by salient stimuli) or endogenously (i.e., driven by one’s goals or intentions). Genetic mouse models are useful tools to investigate the neurobiology of cognition, but a well-established assessment of attention orienting in mice is missing. This study aimed to adapt the Posner task, a widely used attention orienting task in humans, for use in mice using touchscreen technology and to test the effects of two attention-modulating drugs, methylphenidate (MPH) and atomoxetine (ATX), on the performance of mice during this task. In accordance with human performance, mice responded more quickly and more accurately to validly cued targets compared to invalidly cued targets, thus supporting mice as a valid animal model to study the neural mechanisms of attention orienting. This is the first evidence that mice can be trained to voluntarily maintain their nose-poke on a touchscreen and to complete attention orienting tasks using exogenous peripheral cues and endogenous symbolic cues. The results also showed no significant effects of MPH and ATX on attention orienting, although MPH improved overall response times in mice during the exogenous orienting task. In summary, the current study provides a critical translational task for assessing attention orienting in mice and to investigate the effects of attention-modulating drugs on attention orienting.

## Introduction

A fundamental role of attention is to direct an individual’s focus to relevant information in the environment. The ability to selectively attend to a location or modality is referred to as attention orienting [1]. Two types of attention orienting—exogenous and endogenous—have been proposed [1–3]. Exogenous orienting is a stimulus-driven process in which one’s attention is drawn automatically to salient external stimuli. Endogenous orienting represents a goal-directed process in which existing expectations and/or knowledge determine where one’s attention is given. Atypical attention orienting has been found in some individuals with autism spectrum disorder (ASD [4]), social anxiety disorder [5], attention-deficit/hyperactivity disorder (ADHD [6]) and Parkinson’s disease [7]. Limited treatments exist for the orienting deficits in these conditions, largely due to inadequate understanding of the underlying neurobiology of atypical attention orienting.

In humans, exogenous and endogenous orienting are commonly measured by a computer-based visual spatial orienting task designed by Posner and colleagues [8]. In this task, participants are instructed to respond to a left- or right-sided target after the presentation of a cue [1,9]. The exogenous task frequently uses peripheral cues, such as a flash, that are not informative of the target location, whereas the endogenous task typically uses central informative cues, such as arrows, that indicate where the target will appear. The target appears in the cued location during valid trials and in the non-cued location during invalid trials. The outcome measures are reaction time and accuracy. The difference in performance between the valid and invalid trials is referred to as the orienting or validity effect. This effect represents the costs of disengaging and shifting attention from the incorrect to the correct location. If participants are quicker and more accurate at localising targets during valid compared with invalid trials, then their response is regarded as attention orienting that was induced by the perceived cue direction.

The Posner task has been used to investigate the neural basis of attention orienting through neuroimaging and clinical studies [3,10,11]. To allow the study of neural circuits through lesion and pharmacological manipulations, animal models of the Posner task have been developed in recent years. Studies in monkeys and rats showed that attention orienting was affected by lesions of the basal cholinergic nuclei [12] and administration of cholinergic medications [13–16]. Together with clinical findings showing impairments in Alzheimer’s disease, a condition associated with markedly depleted cortical cholinergic innervation [17,18], and beneficial effects of nicotine [19,20], these lines of evidence support acetylcholine (ACh) as a primary neurotransmitter that mediates attention orienting. There is also evidence for the involvement of noradrenaline (NA) and dopamine (DA) in attention orienting in primates [21–23]. NA- and DA-mediating medications, such as methylphenidate (MPH) and atomoxetine (ATX), however, have shown inconsistent effects on attention orienting between humans and non-human primates [23,24]. In contrast to cholinergic involvement, the effect of NA and DA on attention orienting is not well understood.

Genetic mouse models are useful in studying the neural mechanisms underlying cognitive processes [25,26], but a well-established assessment of attention orienting in mice is missing. Recently, Wang and Krauzlis [27] provided the first adaptation of the Posner cueing task in mice. Their study demonstrated that mice exhibited shorter reaction times and higher accuracy to validly cued spatial cues, thus supporting the possibility to measure attention orienting experimentally in mice. These mice, however, were head-fixed, which might have induced high stress that affected attention processes in mice [28]. In addition, the task used peripheral cues to predict the target, which raises the question as to whether mice can endogenously orient their attention based on rule-based symbolic cues like those used in the human Posner task.

The current study aimed to adapt the Posner cueing task for use in mice using touchscreen technology, an increasingly popular method to assess cognitive functions in rodents in a manner that is similar to cognitive tests in humans [29,30]. A major challenge for successful task design is the requirement for mice to be trained to maintain their nose-poke at the touchscreen until the appearance of the target. This is critical to control the distance between the mice and the presentation of the stimuli, to reduce the effects of head movement on vision, and to record an accurate response time. It was hypothesised that mice would respond more quickly and accurately during valid compared with invalid trials in both the exogenous and endogenous task. To further understand the role of the noradrenergic and dopaminergic neurotramitter systems in attention orienting, the study also aimed to explore the effects of clinically effective treatments, methylphenidate (MPH) and atomoxetine (ATX), on mice during the novel Posner-style cueing task. Given that the effects of MPH and ATX on attention orienting were inconsistent in previous research [23,24] and that this study used a novel task, no hypothesis was proposed.

## Materials and Methods

### Animals

Thirty-two male C57BL/6J mice were obtained from the Animal Resources Centre (Murdoch, Western Australia) after weaning at 4 weeks of age. Mice were housed in groups of 4 in individually ventilated cages (39 × 20 × 16 cm) with food and water available *ad libitum*, with shelter, and tissue for nesting. Temperature and humidity were controlled at 22 °C and 45%, respectively. Mice were maintained on a 12-h light/dark cycle (lights on at 0700 hours) and bedding changed weekly. At 7 weeks of age, mice were moved to open-top standard mouse cages (34 × 16 × 16 cm) and to a reversed light cycle (12:12-h, lights off at 0800 hours). Housing groups and shelters were transferred together, though four mice were housed individually to avoid fighting. At 8 weeks of age, mice were weighed daily for three days to determine the baseline free feeding weight (FFW) and then food restricted to 85% FFW. Mice were fed standard chow inside their cages, at the same time of day, with a maximum 0.2 g difference in food weight between days. All procedures were approved by the Florey Institute of Neuroscience and Mental Health Animal Ethics Committee and complied with the relevant guidelines and regulations of the National Health and Medical Research Council Code of Practice for the Use of Animals for Scientific Purposes.

### Drugs and treatments

Methylphenidate (MPH, Cat # M325880, Toronto Research Chemicals) and Atomoxetine (ATX, Cat # Y0001586; Sigma-Aldrich) were dissolved in 0.9% (w/v) sodium chloride, administered intraperitoneally, 30 minutes prior to testing in an injection volume of 10 ml/kg and dose of 3 mg/kg. The doses of MPH and ATX were selected based on previous touchscreen studies that assessed attention in C57BL/6J mice [31].

### Behavioural apparatus

Behavioural testing was conducted in a touchscreen automated operant chamber system (Fig 1a; Campden Instruments Ltd., UK). A black Perspex mask with three square windows (7 × 7 cm, 1.5 cm above the grid floor) was used to cover the touchscreen to reduce incidental touches. Details of the apparatus methods have been described previously [29,32]. Whisker Server and ABET II were used to control the system and to collect data (Lafayette Instruments, Lafayette, IN, USA).

**Figure 1.**
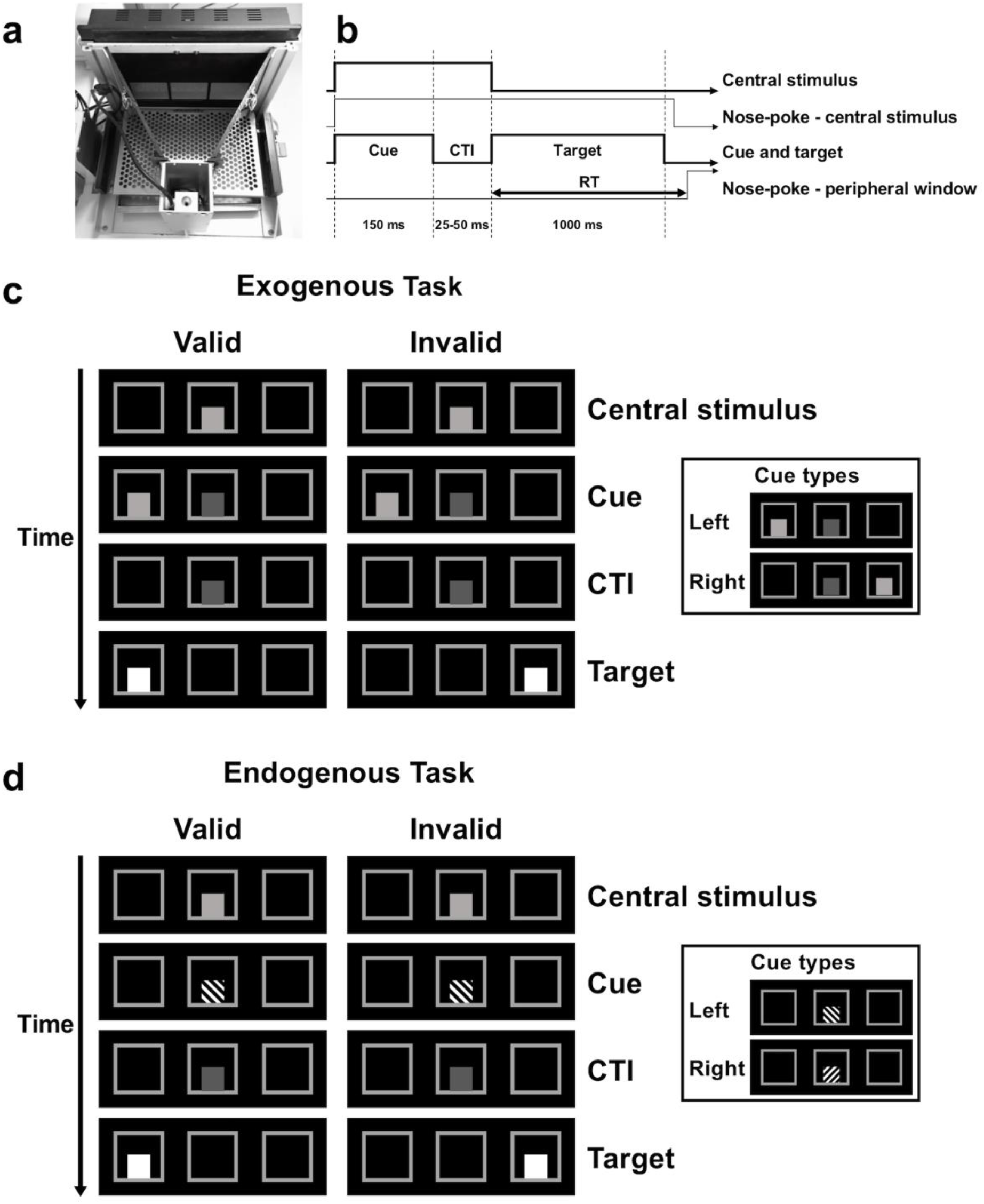
Illustration of the exogenous and endogenous tasks in mice. (a) Photograph of the touchscreen operant chamber. The chamber was equipped with a touchscreen on one end and a liquid-reward (milkshake) delivery magazine on the other. The screen was covered with a black Perspex mask with three response windows. (b) Timing of events in the probe. Trials started with illumination of the central stimulus. After a mouse nose-poked the central stimulus, a cue appeared, followed by the bright peripheral target after a random interval. CTI = cue-target interval. RT = response time. (c) Stimuli in the exogenous task. Cue validity = 50% in the probe. (d) Stimuli in the endogenous task. Cue validity = 80% in the probe.

### Stimuli

Mice were trained to sustain their nose-poke at a stimulus in the central window during the presentation of a cue and then to respond to the target displayed in the left or right window (Fig 1b). All stimuli measured 3.5 × 3.5 cm. The central stimulus was a square at 70% brightness and was dimmed to 20% brightness after being touched by the mouse. The cue for the exogenous task was a square at 70% brightness presented in the peripheral window. The cue for the endogenous task was a square with either 145-degree or 45-degree black grating presented in the central window (Fig 1c & d).

### Pretraining

Mice were habituated to the touchscreen chambers over two 20-min sessions. Following habituation, mice were trained to associate nose-poking at a central stimulus on the touchscreen with a food reward (Iced Strawberry Milk, Nippy’s Ltd, Australia) over two training stages (see Supplementary Materials and Methods).

### Task Training

After completing the pretraining stage, mice were randomly assigned to either the exogenous (*n* = 16) or endogenous task (*n* = 16). The main aim of training was for the mice to nose-poke the central stimulus for the time it took for the cue and the target to be presented. If completed correctly, mice were then rewarded with food delivery. A 5-second inter-trial-interval (ITI) would then elapse before the commencement of the next trial. If mice withdrew their nose from the central stimulus before the onset of the target (anticipation error), a 5-second ITI was initiated with no food reward. Touches to the opposite side of the target (commission error), or failure to respond to the target within a certain time (omission error), would result in no food reward and a 5-second time-out period with illumination of the house light, followed by a 5-second ITI. Omission errors occurred when mice exceeded either the maximum reaction time (i.e., time between target onset and mice leaving the central stimulus) or the maximum movement time (i.e., time between mice leaving the central stimulus and touching the peripheral window).

Each daily training session lasted either 60 minutes or 120 trials (excluding anticipation errors)-whichever came sooner. For both the exogenous and endogenous groups, training was identical except that different cues were used and that mice underwent 6 stages of training for the exogenous task and 7 stages for the endogenous task (Fig 2). For the exogenous task, cue validity was set at 50% in all training stages to prevent mice from learning the association between the cue and the target. For the endogenous task, cue validity was set at 100% in training to facilitate learning that the 145-degree and 45-degree grating cue predicted the left and right target, respectively. In the final training stage for this task, cue validity was reduced to 90% to introduce the invalid endogenous cues to the mice.

**Figure 2.**
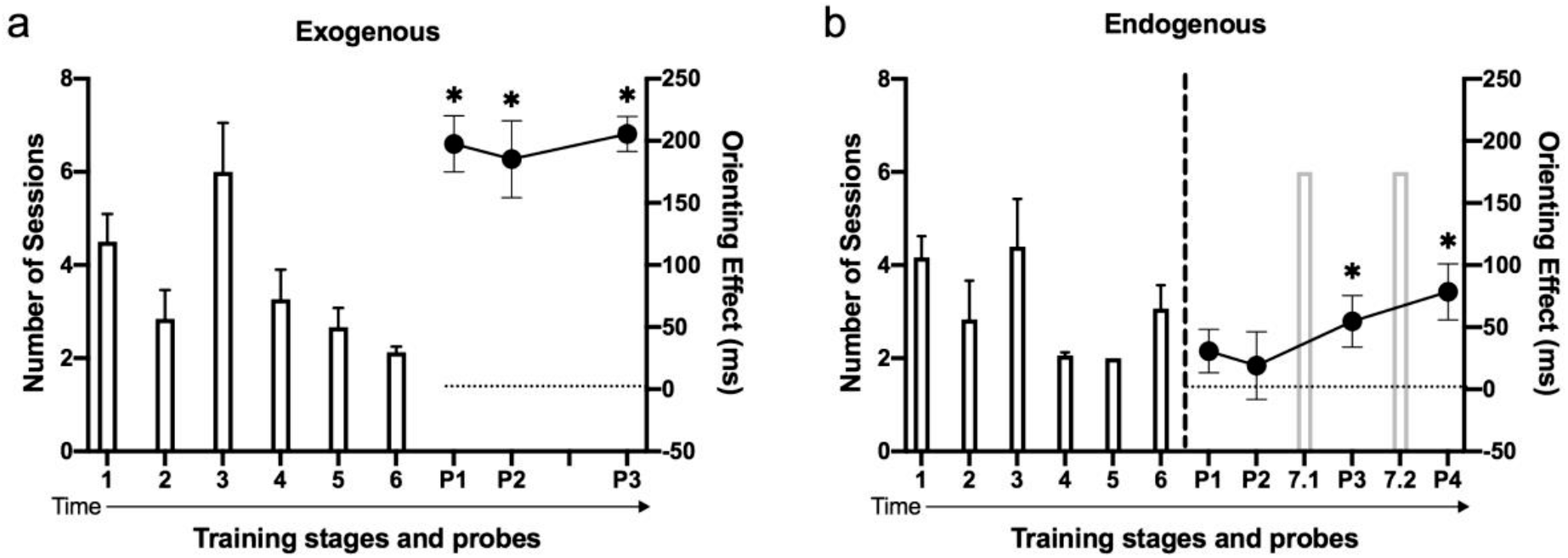
Timeline of training and probes. (a) Exogenous task training. 1 to 6 on the x axis indicate different training stages. P on the x axis indicates the probe. Each probe comprised two sessions. Cue validity was set at 50% in all training sessions. (b) Endogenous task training. Cue validity was 100% in the training stages 1 to 6, 90% in the training stages 7.1 and 7.2, and 80% during the probes. The vertical dashed line denotes the first time that invalid trials were introduced into the endogenous task. Exogenous task: *n* = 16 mice in all training stages; *n* = 9 mice in Probe 1; *n* = 12 mice in Probe 2 and 3. Endogenous task: *n* = 16 mice in all training stages, *n* = 15 mice in all probes. Numbers of sessions to complete each training stage are expressed as bars with mean ± standard errors. Orienting effects in each probe are expressed as circle symbols with mean ± standard errors. * denotes*p* < .05.

For the first three training stages, mice were subjected to stepwise training, in which the duration of cues and cue-target-intervals (CTIs) were adjusted in steps of 50 ms based on the performance of mice. After completing stepwise training, mice were moved to randomised training, in which the cue duration remained at 150 ms and the CTI was randomised between 25 ms and 50 ms. In the last training stage, the target duration was set at 1 s, maximum reaction time at 1.5 s, and the maximum movement time at 2.5 s (see Supplementary Materials and Methods).

### Probes

Mice exhibiting orienting effects [(median RTs in invalid trials – valid trials) > 0] were deemed to be performing the task (Fig 2). In the probes, cue validity remained at 50% in the exogenous task but changed to 80% in the endogenous task.

After mice showed stable orienting, the effects of attention-modulating drugs, MPH and ATX, were assessed. MPH, ATX, and saline were administered in a pseudo-randomised cross-over design with a 3- to 4-day washout period between each administration. Mice were subjected to a minimum of 2 consecutive days of baseline training to ensure continued stable performance between each probe. Mice not reaching criteria (> 70% accuracy or completion of 120 trials, excluding anticipations errors) were subjected to further baseline sessions until criteria were met.

### Data analysis

All data were analysed using generalised linear, latent, and mixed models (GLLAMM) with robust standard error estimation, as previously described [33]. All statistical analyses were performed in STATA (StataCorp, College Station, TX, USA). Graphs were produced using Prism (GraphPad, La Jolla, CA, USA).

## Results

### Mice successfully learnt the exogenous and endogenous tasks

All mice acquired stepwise and randomised training for the exogenous and endogenous tasks (Fig 2; statistical comparisons in Supplementary Results).

### Mice responded more quickly and more accurately to validly cued trials in the exogenous and endogenous tasks

#### Exogenous task

Mice showed longer RTs during invalid relative to valid trials in the last probe (Fig 2a & 3a; Probe 3: coefficient = 198.66, 95% CI = [170.08, 227.25], *p* < .001). The mean orienting effect was 210 ms (Fig 3b; *SE* = 56). Performance on invalid trials also worsened compared to valid trials, with mice showing lower odds of a correct response (Fig 3c; OR = 0.56, 95% CI = [0.42, 0.75],*p* < .001) and higher odds of committing a commission error (Fig 3d; OR = 6.97, 95% CI = [2.94, 16.53], *p* < .001) during invalid trials. There was no significant association between omission errors and cue validity (Fig 3e; OR = 1.31, 95% CI = [0.95, 1.8], *p* = .1). There were no significant interactions between cue validity and nose-poking time or between cue validity and days on RT and on any types of responses, suggesting that the orienting effect was not affected by nose-poking time or days.

**Figure 3.**
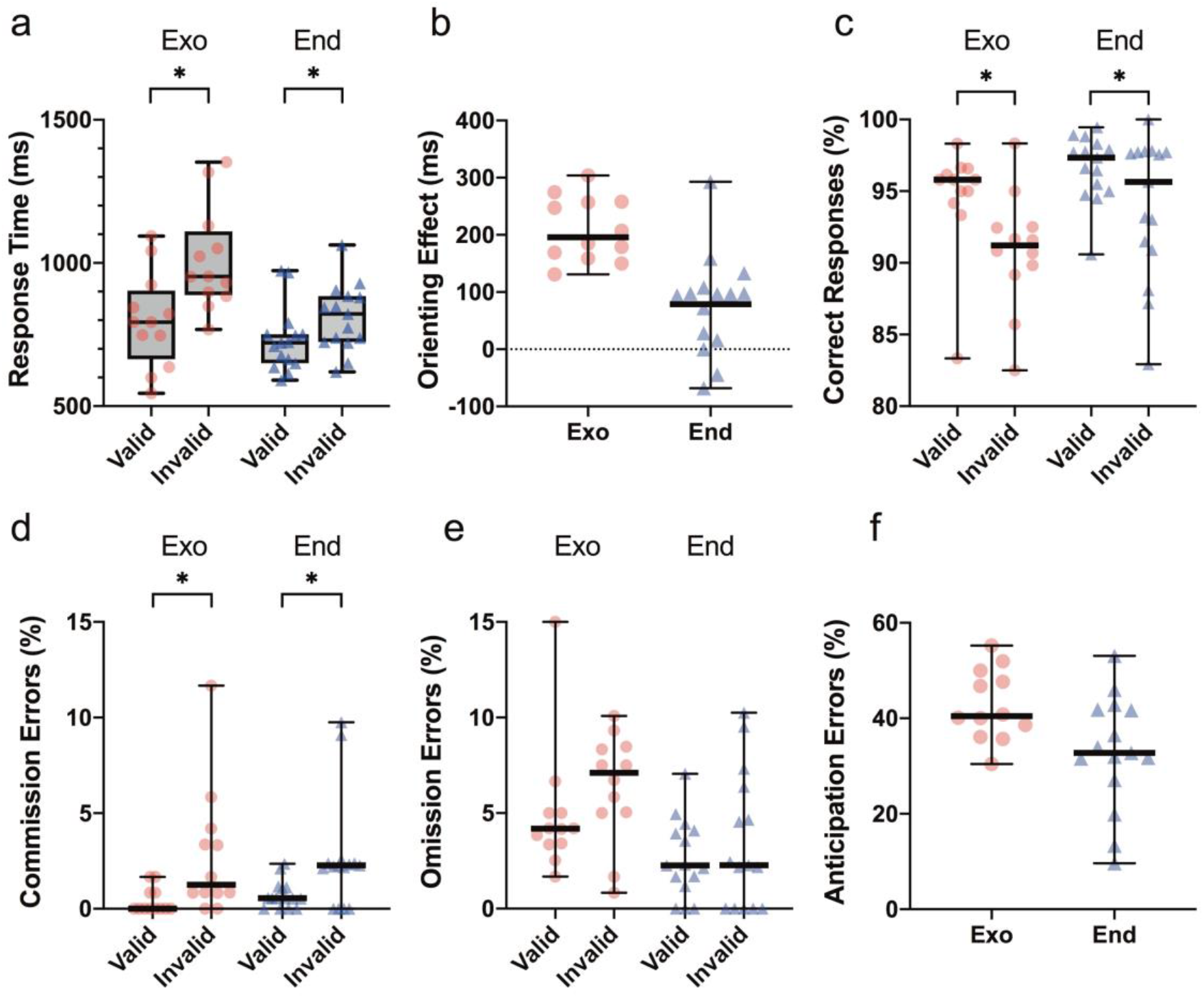
Performance of mice following the completion of training in the exogenous task (Probe 3) and endogenous task (Probe 4). (a) Response time (b) Orienting effect = median RT in invalid trials – median RT in valid trials (c) Correct responses (d) Commission errors (e) Omission errors (f) Anticipation errors. Data in (a) is expressed as box plots with box showing 25th-75th percentile, whiskers showing min-max values, and the central line showing the median. (b-f) are expressed as scatter plots with the long horizontal line showing the median. * denotes *p* < .05. Exogenous *n* = 12 mice; Endogenous *n* = 15 mice.

On average, 43% of the trials initiated by mice were anticipation errors (Fig 3f; *SE* = 5%) in the exogenous task. Mice showed higher odds of an anticipation error when a longer nose-poking time was required (OR = 1.03, 95% CI = [1.03, 1.04], *p* < .001). While mice with higher levels of anticipation errors responded slightly slower to the target overall (coefficient = 16.21, 95% CI = [11.85, 20.58], *p* < .001), no significant effect of anticipation error on the orienting effect (coefficient = 4.04, 95% CI = [−0.26, 8.33], *p* = .07) or any other types of responses was shown.

#### Endogenous task

Mice exhibited longer RTs during invalid compared with valid trials in the last probe (Fig 2a & 3a; Probe 4: coefficient = 79.42, 95% CI = [32.77, 126.08], *p* = .001). The mean orienting effect was 79 ms (Fig. 3b. *SE* = 23). Akin to performance in the exogenous task, mice also showed lower odds of making a correct response (Fig 3c; OR = 0.54, 95% CI = [0.36, 0.79], *p* = .002) and higher odds of committing a commission error (Fig 3d; OR = 3.57, 95% CI = [1.86, 6.87],*p* < .001) during invalid compared with valid trials. There was no association between omission errors and cue validity (Fig 3e; OR = 1.32, 95% CI = [0.8, 2.16], *p* = .28). There were no significant interactions between cue validity and nose-poking time or between cue validity and days on RT and on any types of responses, suggesting that the orienting effect was not affected by nose-poking time or days.

On average, 33% of the trials initiated by mice were anticipation errors (Fig 3f; *SE* = 12%) in the endogenous orienting task. Mice showed higher odds of anticipation errors on trials requiring longer nose-poking time (OR = 1.04, 95% CI = [1.03, 1.04], *p* < .001). In contrast to the exogenous task, higher anticipation errors corresponded to significantly quicker response latencies to the target (coefficient = −4.71, 95% CI = [−8.46, −0.96], *p* = .01). Apart from this, no significant effect of anticipation error on the orienting effect (coefficient = −1.33, 95% CI = [−5.62, 2.96], *p* = .54) or any other types of responses was observed.

### Methylphenidate quickened response time in the exogenous task, while atomoxetine had the opposite effect

During the exogenous task, administration of methylphenidate (MPH) speeded RTs (Fig 4a; coefficient = −119.60, 95% CI = [−118.08, −51.12], *p* = .001) and lowered the odds of omission errors in mice (Fig 4e; OR = 0.74, 95% CI = [0.54, 1], *p* = .049), compared to saline-treated mice. MPH, however, did not alter the orienting effect (Fig 4b; coefficient = 12.9, 95% CI = [−25.6, 51.41], *p* = .51) or the odds of correct responses (Fig 4c; OR = 1.03, 95% CI = [0.79, 1.33], *p* = .83). There was no significant interaction between MPH and cue validity on any outcome measures. MPH appeared to influence measures related to impulsivity, with MPH-treated mice exhibiting higher odds of committing a commission error (Fig 4d; OR = 2.39, 95% CI = [1.38, 4.14], *p* = .002) and higher odds of making an anticipation error (Fig 4f; OR = 1.17, 95% CI = [1.07, 1.27], *p* = .001). The percentage of anticipation errors did not alter the effect of MPH on the outcome measures, except the odds of a correct response being made (OR = 1.03, 95% CI = [1.01, 1.05], *p* = .01).

**Figure 4.**
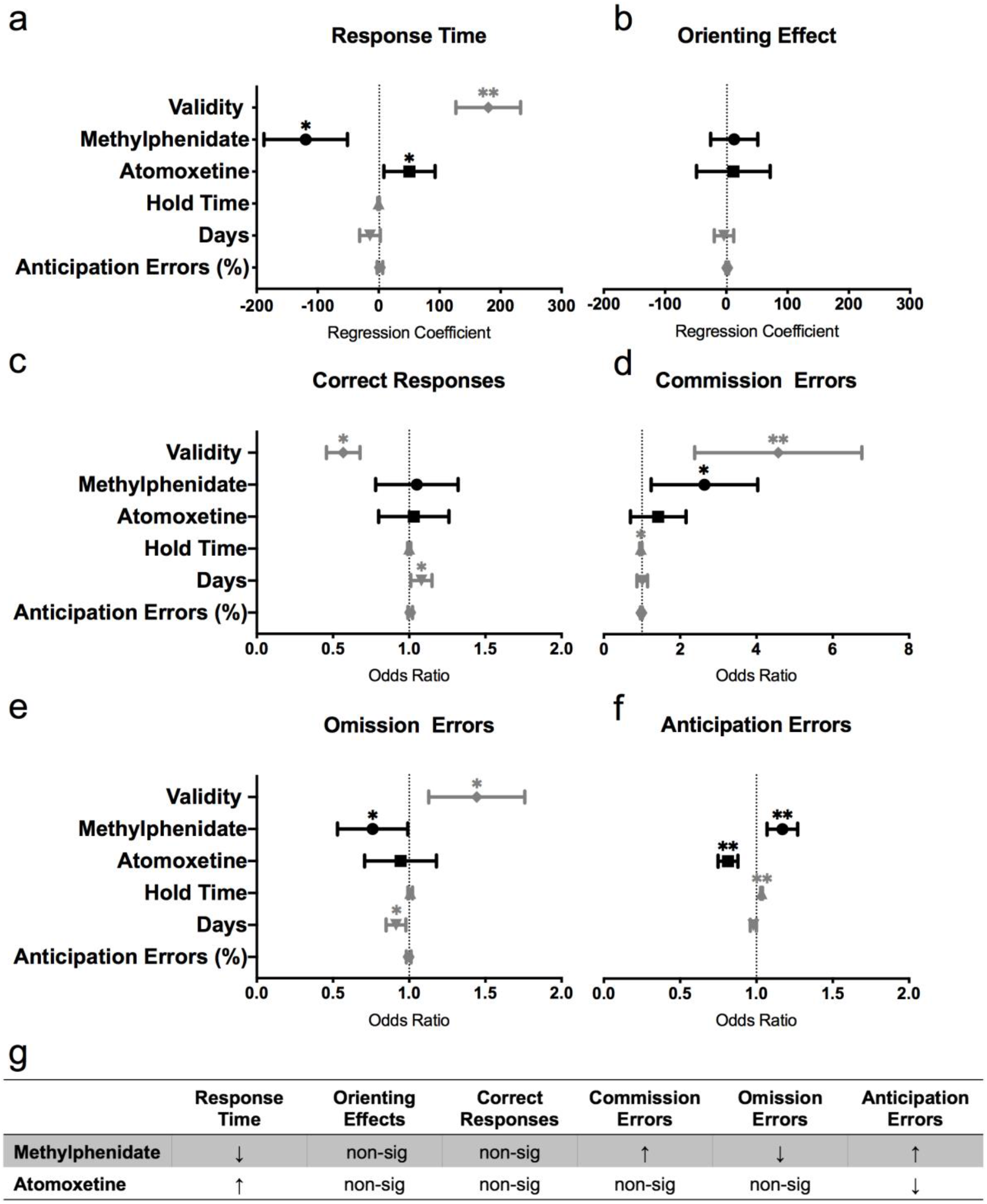
Effects of atomoxetine and methylphenidate on the exogenous task. (a) Response time (b) Orienting effect = median RT in invalid trials – median RT in valid trials (c) Correct responses (d) Commission errors (e) Omission errors (f) Anticipation errors (g) Summary table of the effects of MPH and ATX on the outcome measures. (a) and (b) depict regression coefficients ± 95% confidence intervals. (c) to (f) depict odds ratios ± 95% confidence intervals. * denotes*p* < .05. ** denotes*p* < .001. Saline: *n* = 13 mice; MPH: *n* = 11 mice; ATX: *n* = 13 mice.

Opposite to the effects of MPH, atomoxetine (ATX) slowed RTs in mice compared to saline-treated mice (Fig 4a; coefficient = 50.53, 95% CI = [8.55, 92.51], *p* = .02). ATX did not significantly affect the orienting effect (Fig 4b; coefficient = 11.58, 95% CI = [−48.5, 71.65], *p* = .71), the odds of correct responses (Fig 4c; OR = 1.01, 95% CI = [0.81, 1.27], *p* = .92; Figure 4e), the odds of commission errors (Fig 4d; OR = 1.30, 95% CI = [0.77, 2.21], *p* = .33), or the odds of omission errors (Fig 4e; OR = 0.92, 95% CI = [0.72, 1.19], *p* = .53). There was no significant interaction between ATX and cue validity on any outcome measures. Mice administered ATX showed significantly lower odds of anticipation errors (Fig 4f; OR = 0.81, 95% CI = [0.75, 0.88], *p* = .1), compared to mice on saline. The percentage of anticipation errors did not alter the effect of ATX on the outcome measures.

### Methylphenidate had minimal effect on endogenous task performance, while atomoxetine decreased performance

During the endogenous task, MPH had minimal effect on performance measures. MPH treatment did not affect RTs (Fig 5a; coefficient = 15.85, 95% CI = [−34.71, 66.40], *p* = .54), orienting effects (Fig 5b; coefficient = 24.70, 95% CI = [0.64, 1.29], *p* = .59), the odds of correct responses (Fig 5c; OR = 0.91, 95% CI = [0.79, 1.33], *p* = .83), the odds of commission errors (Fig 5d; OR = 1.94, 95% CI = [0.95, 3.97], *p* = .07), or the odds of omission errors (Fig 5e; OR = 0.93, 95% CI = [0.63, 1.39], *p* = .73) in mice. There was also no significant interaction between MPH and cue validity on any outcome measure. Mice administered with MPH did show higher odds of committing an anticipation error (Fig 5f; OR = 1.49, 95% CI = [1.36, 1.64],*p* < .001), compared to mice on saline. The percentage of anticipation errors did not alter the effect of MPH on the outcome measures, except the odds of commission errors (OR = 0.94, 95% CI = [0.89, 0.99], *p* = .003).

ATX impaired endogenous task performance in mice, across most of the measures. Mice administered with ATX showed significantly slower RTs (Fig 5a; coefficient = 39.42, 95% CI = [14.56, 64.29], *p* = .002;), lower odds of correct responses (Fig 5c; OR = 0.72, 95% CI = [0.54, 0.97], *p* = .03), higher odds of omission errors (Fig 5e; OR = 1.44, 95% CI = [1.05, 1.98], *p* = .03), and higher odds of anticipation errors (Fig 5f; OR = 1.18, 95% CI = [1.08, 1.29], *p* < .001). ATX did not significantly affect orienting effects (Fig 5b; coefficient = 22.52, 95% CI = [−22.55, 67.59], *p* = .33) or the odds of commission errors (Fig 5d; OR = 1.11, 95% CI = [0.51, 2.40], *p* = .26). There was no significant interaction between ATX and cue validity on any outcome measure. The percentage of anticipation errors did not alter the effect of ATX on the outcome measures, except the odds of correct response (OR = 1.03, 95% CI = [1.00, 1.05], *p* = .004).

**Figure 5.**
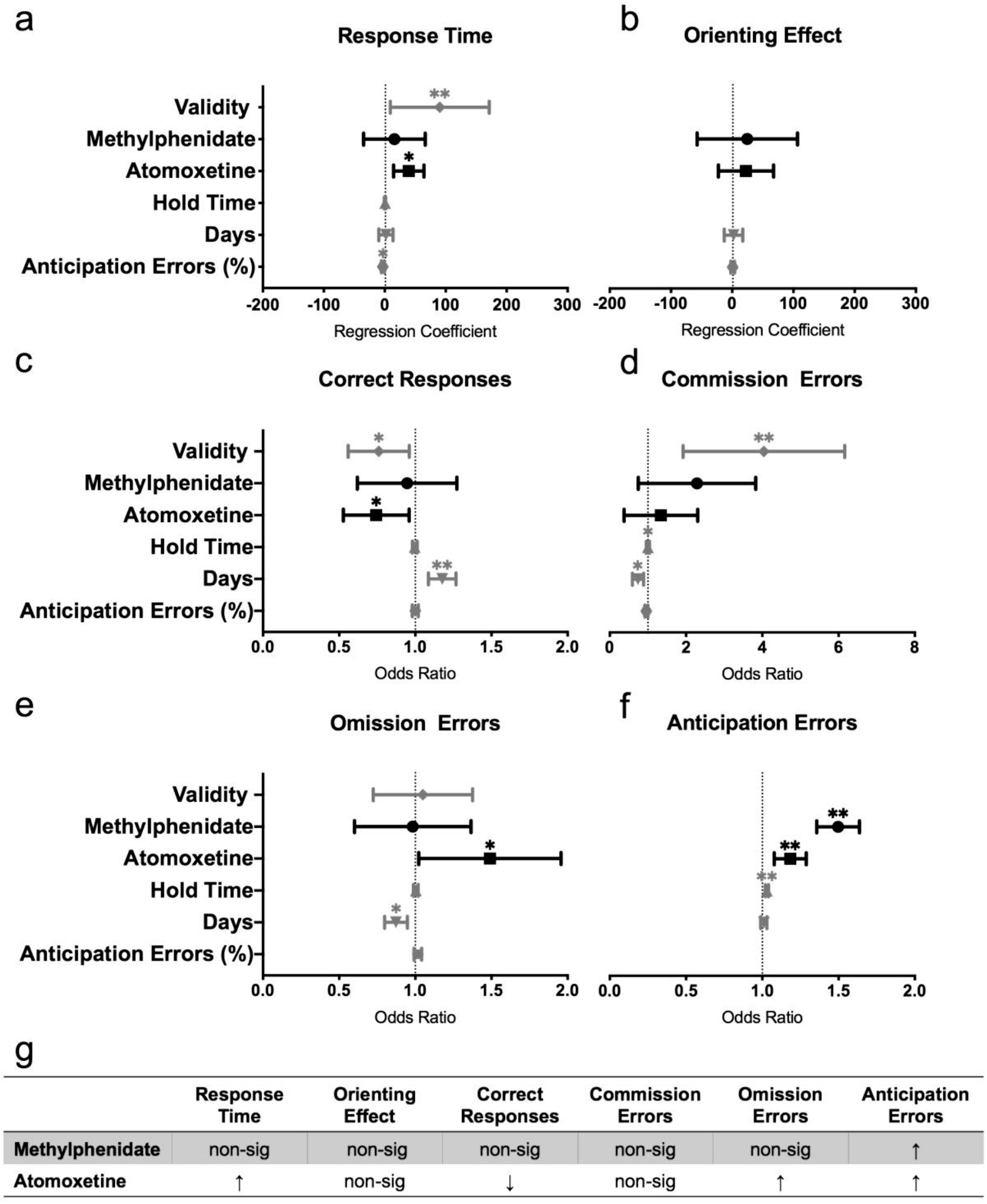
Effects of atomoxetine and methylphenidate on the endogenous task. (a) Response time (b) Orienting effect = median RT in invalid trials – median RT in valid trials (c) Correct responses (d) Commission errors (e) Omission errors (f) Anticipation errors (g) Summary table of the effects of MPH and ATX on the outcome measures. (a) and (b) depict regression coefficients ± 95% confidence intervals. (c) to (f) depict odds ratios ± 95% confidence intervals. * denotes*p* < .05. ** denotes*p* < .001. Saline: *n* = 15 mice; MPH: *n* = 14 mice; ATX: *n* = 15 mice.

## Discussion

Mice can orient their attention both exogenously and endogenously, as assessed by a new touchscreen-based task adapted from the human Posner task. Like previous results in humans, mice responded more quickly and accurately to validly cued stimuli in both the exogenous and endogenous tasks. During the exogenous task, MPH administration resulted in a more impulsive and alert response style, with a speeding of response times and a reduction in omission errors, but an increase in commission and anticipation errors. ATX administration, in contrast, resulted in a more cautious response style, with a slowing of response times and a lower rate of anticipation errors. During the endogenous task, MPH administration had minimal effect, whereas ATX administration resulted in a decrease in overall performance—slowed RTs, increased odds of omission and anticipation errors, and decreased odds of correct responses being made. Although MPH and ATX showed differential effects on the performance of the mice during the attention orienting tasks, neither treatment altered attention orienting. Overall, the current study provides a novel protocol to investigate the neural mechanisms of attention orienting in mice. Using this new protocol, the current study showed that mice can be trained to voluntarily engage during the attention orienting tasks and, more importantly, can endogenously orient their attention based on learnt symbolic cues.

### Task acquisition: mice can be trained to spontaneously complete the attention orienting tasks

An important achievement in the current study was to train the mice to voluntarily head fix at the centre of the touchscreen until the presentation of the target after a cue, which is critical in the successful adaption of the Posner task to mice. The head fixing behaviour helps to control the starting position of the mice in each trial to obtain reasonably accurate measurements of whole-body response times. The head fixing behaviour was also designed to resemble the procedure of the human Posner task in which participants are required to orient their attention to the appearance of the peripheral targets without moving their eyes or head. As mice lack fovea in their retinas, their orienting is primarily by head and body rather than eye movements [34]. By training the mice to maintain their heads centrally, their eyes were placed to see the central and peripheral stimuli. Previous studies have successfully trained rats to sustain the nose-poke at the centre in the adapted Posner task [16,35]. Similar to rats, the current study showed that mice could be trained to extend their nose-poking time on the touchscreen after stepwise training and to maintain their nose-poking ability during the randomised training. This, to our knowledge, has not been reported in previous research.

### Probe performance: mice showed both stimulus-driven and goal-driven orienting of attention

The results of the current study suggested that mice could orient their attention based on both exogenous and endogenous spatial cues, supporting the use of mice as a valid animal model to study the neural mechanisms of attention orienting. Mice were faster and more accurate at responding to validly cued targets, consistent with findings in humans [8,36] and rats [35]. Similarly, a previous study showed that mice could use spatial cues to orient their attention [27]. In this previous study, however, the heads of the mice were restrained by an implanted head post, which might create stress and is not comparable with the human task. Using touchscreen technology, the current study extends this previous study by showing that mice can spontaneously orient their attention in a low stress setting that is directly compatible with human variations of the tasks.

The current study provides the first demonstration of endogenous orienting in mice based on predictive symbolic cues. Previous rodent studies have typically employed predictive peripheral cues to measure endogenous orienting [35]. Predictive peripheral cues, however, have been suggested to produce both an exogenous attention capture and an endogenous shift of attention [2,37], which may confound the measurement. The current study used symbolic cues (i.e., diagonal stripes) that predicted the location of the targets in the endogenous task, which closely resembles the endogenous cues in the human Posner task [38]. Mice were faster and more accurate when the symbolic cues correctly predicted the location of the targets. This finding opens up avenues for future studies to investigate endogenous orienting in mice using symbolic cues to reduce the confounding components of peripheral cues.

Mice showed a high percentage of anticipation errors (i.e., leaving the touchscreen before the onset of target) in the novel attention orienting task. This finding is different from the performance of rats in a Posner-style cueing task [35]. In that study, the rats made few anticipation errors when the interval between the cue and target was 200 ms, which is similar to the timing used in the current study. It is likely that mice are inherently more active and impulsive, as previous findings also showed that mice tended to respond before stimulus onset in the 5-choice serial reaction time task (5-CSRTT), a task used to measure visual attention and impulsive actions [39,40]. The difference in anticipation errors between mice and rats may also be due to the different apparati used. In the rat study, animals were required to poke their nose into a hole rather than touching a screen, which may have prevented them from quickly withdrawing their nose poke. Although mice showed a high percentage of anticipation errors, the current results demonstrated that the mice completed 120 trials with high accuracy of responses (> 90%), other than anticipation errors, in each task session. In addition, the percentage of anticipation errors in mice did not affect the orienting effect or the odds of making correct responses or errors. Together, these findings suggest that despite the high percentage of anticipation errors, mice were able to complete the attention orienting tasks properly when they could maintain their nose-poke until the appearance of the target.

### Effects of drugs: MPH and ATX exerted mixed effects on exogenous and endogenous orienting in mice

In order to investigate the neural mechanisms of attention orienting, two attention-modulating treatments, methylphenidate (MPH) and atomoxetine (ATX), were administered to the mice in the current tasks. MPH and ATX were observed to improve sustained and selective attention in mice in some studies [31,41], but neither treatment exhibited significant effects on attention orienting (i.e., the orienting effects) in the current tasks. One possible explanation is that the noradrenergic (NA) and dopaminergic (DA) signalling systems, that are modulated by these compounds, have limited or indirect effects on attention orienting. Some studies suggest that NA specifically impacts alertness rather than attention orienting [42,43], although other evidence suggests that NA might affect attention orienting through facilitating the action of acetylcholine (ACh [44]). It has also been suggested that DA is involved in resolving conflict rather than attention orienting [44,45]. Another possible reason is that attention-modulating drugs may only show enhancing effects when a primary attention deficit is present. Previous studies did not find consistent improvement of MPH or ATX on attention in healthy humans and animals [23,31,46,47]. It is possible that healthy subjects might be able to orient their attention near their peak level, and any additional increase of noradrenergic and dopaminergic activities induced by the drugs would not necessarily improve performance. To further investigate this issue, it might be necessary to test mouse models with potential deficits in attention, or use drugs that decrease their attention functioning, such as cholinergic antagonists [15,16].

Although MPH and ATX did not significantly affect attention orienting in mice, these treatments exerted differential effects on behavioural performance. In the exogenous task, mice administered MPH appeared to be more alert, as indicated by faster responses and reduced odds of making omission errors, and more impulsive, as indicated by the increased odds of making commission and anticipation errors. In contrast, mice on ATX tended to be more cautious, as indicated by slower responses and lower odds of anticipation errors. Similarly, previous studies have suggested that ATX reduces impulsive actions in rodents [39,48,49] and humans [50,51], in contrast to MPH [31,52,53]. The differential effects of MPH and ATX on impulsive actions has been suggested to reflect their differential roles on subcortical dopamine neurotransmission [24,41].

During the endogenous task, MPH exerted minimal impact on the performance of mice, with the exception of increasing the odds of anticipation errors. The minimal effect of MPH was partially supported by an early study in healthy people, in which MPH speeded overall RTs but did not affect the orienting effect [54]. In contrast, ATX administration in mice slowed response times, lowered the odds of making a correct response, and increased the odds of making omission and anticipation errors. Due to limited previous research in this area, it is unclear why ATX appeared to impair endogenous orienting. One possible explanation is that although the dose of ATX used in the current study was based on previous research [31], it may lead to an atypical level of arousal for the endogenous orienting task. Endogenous orienting is a goal-directed cognitive process that initially requires mice to maintain a higher level of intrinsic arousal relative to other attention processes. The equivalent dose of ATX that enhanced the performance of mice in other tasks, such as the continuous performance test [31], may have led to a state of hypo- or hyper-arousal in the endogenous orienting task, affecting the ability of the mouse to sustain attention to the task. According to the inverted U-shaped arousal-performance theory, optimal performance occurs at an intermediate level of arousal, whereas high or low levels of arousal will impair performance [55,56]. To understand the pharmacological effects on the endogenous orienting task in mice, future studies are needed to examine the effects of different doses of drugs on task performance.

## Conclusion

This study provides a novel mouse attention orienting task based on the human Posner task. In accordance with human performance, mice responded more quickly and more accurately to validly cued targets, supporting mice as a valid animal model to study the neural mechanisms of attention orienting. Our results provide the first evidence that mice can be trained to voluntarily maintain their nose-poke on the touchscreen and complete both the exogenous and endogenous orienting tasks. These findings support the use of touchscreen to accurately record response time in mice, which has substantial translational relevance due to the reliance on response time measurement in human studies. This study is also the first to show that mice can orient their attention based on the rule-based symbolic endogenous cues, constituting a more accurate measurement of endogenous orienting compared to peripheral cues.

Our results did not show significant effects of MPH and ATX on attention orienting, although MPH improved overall response times in mice during the exogenous orienting task. This is the first study to examine pharmacological effects on attention orienting in mice using our newly developed task. This paves the way for future research to investigate the effects of attention-modulating drugs, and other therapeutic interventions, on attention orienting and to evaluate mouse models of attention disorders.

## Supporting information

Supplementary Results

## Acknowledgments

Thank you to A/Prof. Gilberto Fernando Xavier and Mr. Mateus Torres Cruz for providing advice on the development of the training protocols. Thank you to Mr. Daniel Drieberg for providing animal care. Thank you to Mr. Brett Purcell for providing technical support.

## Author Contributions

SL, KAJ, and ELB conceived and designed the study. SL and ELB carried out the experiments. SL analysed the data with help from CM and EB. SL, KAJ, and ELB contributed to the interpretation of the results. SL wrote the manuscript with input from KAJ, ELB, and AJH. All authors read the final version and consented to publishing this manuscript.

