## Supplementary Materials and Methods for "Assessing Attention Orienting in Mice: A Novel Touchscreen Adaptation of the Posner-Style Cueing Task"

***Pretraining***

Mice were habituated to the touchscreen chambers over two 20-min sessions. During habituation, a food reward (Strawberry Milk, Nippy's Ltd, Australia) was delivered every 10 s. Food reward volume was set at 7 μl in all training and testing stages. Following habituation, mice were trained to associate nose-poking at a central stimulus on the touchscreen with the food reward over two training stages—the Initial Touch stage and the Must Touch stage. In the Initial Touch stage, a food reward was delivered for every 30 s. Three times of the food reward would be delivered if mice touched the central stimulus. In the Must Touch stage, no food reward was given unless the mice touched the central stimulus. The completion criteria was to collect 30 food rewards within 30 mins for the Initial Touch stage, and to collect 60 rewards within 45 mins for the Must Touch stage. It took 1 day for mice to complete the Initial Touch stage, and 1 to 3 days to complete the Must Touch stage.

***Stepwise training***

In these training stages, mice were rewarded for sustaining their nose poke in iterative steps, until they were able to remain still from the onset of the central stimulus, the duration of the cue presentation, and to the onset of the target. In the first three training stages, the cue duration was increased by 50 ms if mice completed 3 consecutive trials (i.e., no anticipation errors). After the cue duration was increased to 150 ms, a 50 ms cue-target-interval (CTI) was introduced. If mice made 5 consecutive anticipation errors, the CTI was removed, after which the cue duration was decreased in steps of 50 ms. The target duration was set at 5 s during stepwise training. The minimum sustained nose-poking times for Training Stages 1, 2, and 3 were 0 ms, 50 ms, and 100 ms, respectively. Mice completed each stepwise training stage when they 1) completed 120 trials (excluding anticipation errors), 2) achieved at least 70% accuracy (correct choice / (total trials, excluding anticipation errors)), and 3) completed no more than 30 trials at the minimum nose-poking time, for two consecutive training sessions.

***Randomised training***

In the randomised training stages, cue duration remained at 150 ms and the CTI was randomised between 25 ms and 50 ms. In Stage 4, 5, and 6, target duration was set at 5 s, 3 s, and 1 s and the maximum reaction time was set at 5.5 s, 3.5 s, and 1.5 s, respectively. The maximum movement time was the same as the maximum reaction time in Stage 4 and 5 and was reduced to 2.5 s in Stage 6. Mice in the endogenous task underwent Stage 7, which was the same as Stage 6 except for the inclusion of 10% invalid trials. The completion criteria for each randomised training stage were 1) completing 120 trials (excluding anticipation errors), and 2) at least 70% accuracy (correct choice / (total trials, excluding anticipation errors)), for two consecutive training sessions.

***Data analysis***

Data were analysed separately for each task. For each trial, responses were recorded as an anticipation error if the mouse was unsuccessful in maintaining their nose-poke to the onset of the target. Otherwise, responses were recorded as either correct response, commission error, or omission error. These responses were treated as binary outcomes and coded yes = 1, no = 0. Response times (RTs, time between the target onset and screen touch) were recorded when mice made correct responses to the target from each trial and were used to calculate orienting effects (median RTs in invalid trials – valid trials) from each session. Both RTs and orienting effects were treated as continuous variables.

All data were analysed using generalised linear, latent, and mixed models (GLLAMM) with robust standard error estimation, as previously described [1]. Individual animals were treated as random effects to reflect the clustered nature of all observations within each animal. GLLAMM were run with cue type (valid or invalid), nose-poking time, testing day, and drug (if appropriate) as independent variables, and with correct responses, commission errors, omission errors, response times, and orienting effects as outcome variables. To determine the effect of anticipation errors on outcome variables, the percentage of anticipation errors for each mouse in each session was calculated and regressed against outcome variables. The adjusted effect of the independent variables on the outcome variables were estimated as odds ratios (ORs) by random-effects logistic regression (binary data) or as median value (coefficients) by median regression (continuous data). The two-tailed *p*-values and the corresponding 95% confidence intervals were reported to indicate the estimates’ precision. No further analyses would be conducted if the interaction effect was trivial (i.e., 1.5^−1^ < OR < 1.5 [2], or median value < 10 ms), irrespective of the statistical significance.

Mice failing to reach baseline criteria and/or to complete 120 trials (excluding anticipation errors) in probe sessions were excluded from data. No mice failed to reach criteria for more than 3 consecutive baseline sessions. Final subject numbers for each group in each probe are reported in the figure legends (Fig 2 to 5). Due to technical issues, a small number of trials were excluded for each mouse in each session. All statistical analyses were performed in STATA (StataCorp, College Station, TX, USA). Graphs were produced using Prism (GraphPad, La Jolla, CA, USA).
