## Supplementary Results for "Assessing Attention Orienting in Mice: A Novel Touchscreen Adaptation of the Posner-Style Cueing Task"

***Mice successfully learnt the exogenous and endogenous tasks***

In the early three training stages, the duration that mice were required to sustain their nose-poke at the centre was adjusted based on the individual performance of each animal, in a stepwise manner, from 0 ms to 200 ms. The mean number of sessions to complete Stages 1-3 for the exogenous task was 6 (*n* = 12, *SE* = 1), 3 (*n* = 13, *SE* = 1), and 6 (*n* = 10, *SE* = 1), respectively (Fig 2a), and for the endogenous task was 4 (*n* = 12, *SE* = 0), 3 (*n* = 12, *SE* = 1), and 4 (*n* = 15, *SE* = 1), respectively (Fig 2b). Following the stepwise training, mice were moved to randomised training, in which cue duration was set at 150 ms, and the cue-target-interval (CTI) was randomised between 25 ms and 50 ms to prevent the mice from developing a habitual response of leaving the touchscreen after a fixed interval. There were three randomised training stages, each with shorter target durations and shorter maximum response time. The mean number of sessions to complete Stages 4-6 for the exogenous task was 3 (*n* = 15, *SE* = 1), 3 (*n* = 15, *SE* = 0), and 2 (*n* = 16, *SE* = 0), respectively (Fig 2a), and for the endogenous task was 2 (*n* = 16, *SE* = 0), 2 (*n* = 16, *SE* = 0), and 3 (*n* = 15, *SE* = 1), respectively (Fig 2b).

To determine whether mice were using the cues to anticipate where the target would appear, the difference in response times (RTs) between the invalidly and validly cued trials (i.e., the orienting effect) was examined. If mice exhibited significantly positive orienting effects, then they were deemed to have learnt the task. Performance was examined in two consecutive probes after the completion of the randomised training. Mice in the exogenous task showed longer RTs in invalid trials compared to valid trials (Fig 2a. Probe 1: coefficient = 192.77, 95% CI = [152.48, 233.05], *p* < .001; Probe 2: coefficient = 171.50, 95% CI = [135.65, 207.35]. The mean orienting effects in Probe 1 and 2 were 198 ms (*SE* = 20) and 190 ms (*SE* = 26), respectively. When performance in the endogenous group was probed, mice did not show a significant difference in RTs between the invalid and valid trials (Fig 2b. Probe 1: coefficient = 26.04, 95% CI = [-15.27, 67.34], *p* = .22; Probe 2: coefficient = 34.62, 95% CI = [-10.37, 79.61], *p* = .13), indicating that they were not using the cues to predict where the target stimuli would appear and thus were not performing the task. We tested the hypothesis that exposing mice to invalid trials in the endogenous task training would improve their performance in the probe. Mice were subjected to 6 days of training with the inclusion of 10% invalid trials (Fig 2b; Training 7.1). This additional training improved performance, with mice showing significantly longer RTs in invalid trials versus valid trials (Fig 2b. Probe 3; coefficient = 56.86, 95% CI = [18.60, 95.12], *p* = .004). The mean orienting effect was 55 ms (*SE* = 21). Mice in the exogenous task were rested for 20 days, with a refresher baseline session every 6 days, while mice in the endogenous task undertook further training (Fig 2b; Training 7.2). Another probe was conducted to examine whether the orienting effect was stable. Results from these final probes are presented in detail in the article.
